# Neuropeptide Modulation Enables Biphasic Inter-network Coordination via a Dual-Network Neuron

**DOI:** 10.1101/2024.03.19.585771

**Authors:** Barathan Gnanabharathi, Savanna-Rae H Fahoum, Dawn M Blitz

**Author notes:** **Correspondence should be addressed to:** Dawn M. Blitz, 700 E High St, PSN 212, Oxford OH, 45056.

## Abstract

Linked rhythmic behaviors, such as respiration/locomotion or swallowing/chewing often require coordination for proper function. Despite its prevalence, the cellular mechanisms controlling coordination of the underlying neural networks remain undetermined in most systems. We use the stomatogastric nervous system of the crab *Cancer borealis* to investigate mechanisms of inter-network coordination, due to its small, well characterized feeding-related networks (gastric mill [chewing, ∼0.1 Hz]; pyloric [filtering food, ∼1 Hz]). Here, we investigate coordination between these networks during the Gly^1^-SIFamide neuropeptide modulatory state. Gly^1^-SIFamide activates a unique triphasic gastric mill rhythm in which the typically pyloric-only LPG neuron generates dual pyloric- plus gastric mill-timed oscillations. Additionally, the pyloric rhythm exhibits shorter cycles during gastric mill rhythm-timed LPG bursts, and longer cycles during IC, or IC plus LG gastric mill neuron bursts. Photoinactivation revealed that LPG is necessary to shorten pyloric cycle period, likely through its rectified electrical coupling to pyloric pacemaker neurons. Hyperpolarizing current injections demonstrated that although LG bursting enables IC bursts, only gastric mill rhythm bursts in IC are necessary to prolong the pyloric cycle period. Surprisingly, LPG photoinactivation also eliminated prolonged pyloric cycles, without changing IC firing frequency or gastric mill burst duration, suggesting that pyloric cycles are prolonged via IC synaptic inhibition of LPG, which indirectly slows the pyloric pacemakers via electrical coupling. Thus, the same dual-network neuron directly conveys excitation from its endogenous bursting and indirectly funnels synaptic inhibition to enable one network to alternately decrease and increase the cycle period of a related network.

**Significance Statement:** Related rhythmic behaviors frequently exhibit coordination, yet the cellular mechanisms coordinating the underlying neural networks are not determined in most systems. We investigated coordination between two small, well-characterized crustacean feeding-associated networks during a neuropeptide-elicited modulatory state. We find that a dual fast/slow network neuron directly shortens fast network cycles during its slow, intrinsically generated bursts, likely via electrical coupling to fast network pacemakers, despite rectification favoring the opposite direction. Additionally, the fast network is indirectly prolonged during another slow-network phase, via chemical synaptic inhibition that is likely funneled through the same electrical synapse. Thus, a dual-network neuron alternately reinforces and diminishes neuropeptide actions, enabling distinct frequencies of a faster network across different phases of a related slower rhythm.

## Introduction

Many rhythmic behaviors require coordination for proper function, such as locomotion and respiration, and multiple orofacial behaviors (e.g., respiration, swallowing, chewing, vocalization) (Barnett et al., 2021; Cao et al., 2012; Hao et al., 2021; Huff et al., 2022; Juvin et al., 2022; Wei et al., 2022). Central pattern generator (CPG) networks controlling rhythmic behaviors are flexible to accommodate changing organismal needs (Bucher et al., 2015; Grillner and El Manira, 2020; Ramirez and Baertsch, 2018). Behavioral coordination requires coordinating the underlying CPG networks. Significant consequences occur from disrupted coordination, such as aspiration of food if swallowing and respiration are not coordinated (Yagi et al., 2017). Thus, it is important to understand the cellular-level mechanisms controlling inter-network coordination.

Network coordination can occur as correlated changes in frequency or strength, as for respiration with changing locomotor speeds or at the onset of vocalizations, or as specific timing relationships to coordinate movements of different body parts (Anderson et al., 2016; Daley et al., 2013; Demartsev et al., 2022; Juvin et al., 2022; Mulloney and Smarandache-Wellmann, 2012; Ramirez and Baertsch, 2018; Wei et al., 2022). Such interactions may occur via direct CPG to CPG projections, or indirectly via feedback from networks to higher-order inputs to a related CPG (Bartos and Nusbaum, 1997; Gariépy et al., 2010; Wood et al., 2004). Additional anatomical substrates for network coordination include sensory feedback from one behavior influencing related CPGs, or parallel descending input to related networks (Barnett et al., 2021; Gariépy et al., 2010; Juvin et al., 2022). More complex behavioral interactions may also occur, such as variations in respiration during crying, singing, or while playing wind instruments (Fortune et al., 2011; Hérent et al., 2023; Higashino et al., 2022; Okobi et al., 2019; Wei et al., 2022). Further, coordination can be altered by developmental, physiological, or environmental stimuli (Blitz and Nusbaum, 2008; Clemens et al., 1998a; Moore et al., 2013; Saunders et al., 2004; Sillar et al., 2023; Stein and Harzsch, 2021). Although insights have been gained into inter-network coordination, its complexity and plasticity coupled with often large, distributed networks make it difficult to fully determine how network coordination is controlled (Juvin et al., 2022).

Here we used small, well-described networks in the stomatogastric nervous system (STNS) of the Jonah crab, *Cancer borealis*, to investigate cellular-level mechanisms of inter-network coordination (Bartos et al., 1999; Bartos and Nusbaum, 1997; Daur et al., 2016; Wood et al., 2004). An in vitro STNS preparation includes the stomatogastric ganglion (STG, ∼30 neurons) containing well-characterized CPGs that control the gastric mill (food chewing) and pyloric (food filtering) rhythms, plus identified modulatory and sensory inputs (Fig. 1) (Blitz, 2023; Daur et al., 2016; Nusbaum and Beenhakker, 2002). The pacemaker-driven pyloric rhythm is constitutively active in vivo and in vitro (∼1 s cycle period), whereas the slower (∼10 s cycle period) network-driven gastric mill CPG requires activation by modulatory or sensory inputs (Blitz, 2023; Nusbaum and Beenhakker, 2002). All pyloric and gastric mill neurons (1-5 neurons/type) and their transmitters are identified, and their connectivity is mapped under multiple modulatory conditions (Fig. 1B) (Daur et al., 2016; Fahoum and Blitz, 2024; Marder and Bucher, 2007). Although the gastric mill and pyloric rhythms can occur independently, they are often coordinated, and the extent and type of coordination is altered by modulatory, behavioral, and environmental conditions (Bartos et al., 1999; Bartos and Nusbaum, 1997; Blitz et al., 2019; Clemens et al., 1998a; Stein and Harzsch, 2021; Wood et al., 2004).

We investigated mechanisms of coordination during the modulatory state elicited by the modulatory commissural neuron 5 (MCN5), or bath application of its neuropeptide Gly^1^-SIFamide (Blitz et al., 2019; Fahoum and Blitz, 2021; Fahoum and Blitz, 2024). In this state, the typically pyloric-only LPG switches into dual pyloric and gastric mill rhythm participation, becoming the third phase of a gastric mill rhythm (Fahoum and Blitz, 2024). This contrasts with biphasic gastric mill rhythm versions (Beenhakker et al., 2004; Blitz et al., 2008, 1999; Christie et al., 2004). The pyloric rhythm cycle period varies across the Gly^1^-SIFamide gastric mill rhythm, but a full description plus identification of mechanisms of interactions between the two networks is lacking (Blitz et al., 2019). The well-described connectome and small populations enabled selective manipulation of neuronal populations to determine the full extent of gastric mill regulation of the pyloric rhythm and its underlying cellular mechanisms during the unique triphasic Gly^1^-SIFamide-elicited gastric mill rhythm. Our results highlight the complexity of synaptic connectivity that can mediate coordination between related networks.

## Methods

### Animals

Wild caught adult male *C. borealis* crabs were procured from The Fresh Lobster Company (Gloucester, MA), maintained in artificial seawater (10-12°C) tanks, and fed twice weekly until used. Prior to dissection, crabs were anesthetized by cold packing in ice for 40 – 50 minutes. The crab foregut was first removed, bisected, and pinned flat, ventral side up, in a Sylgard 170-lined dish (Thermo Fisher Scientific). The STNS was then dissected free from muscles and connective tissue and pinned in a Sylgard 184-lined petri dish (Thermo Fisher Scientific) (Fahoum and Blitz, 2021; Gutierrez and Grashow, 2009). The preparation was kept in chilled (4°C) *C. borealis* physiological saline throughout dissections.

### Solutions

*C. borealis* physiological saline was composed of 440 mM NaCl, 26 mM MgCl_2_, 13 mM CaCl_2_, 11 mM KCl, 10 mM Trizma base, 5 mM maleic acid (pH 7.4-7.6). Squid internal electrode solution contained 10 mM MgCl_2_, 400 mM potassium D-gluconic acid, 10 mM HEPES, 15 mM Na_2_SO_4_, 20 mM NaCl, pH 7.45 (Hooper et al., 2015). Gly^1^-SIFamide (GYRKPPFNG-SIFamide, custom peptide synthesis: Genscript) (Blitz et al., 2019; Dickinson et al., 2008; Huybrechts et al., 2003; Yasuda et al., 2004) was prepared by dissolving it in optima water (Thermo Fisher Scientific) at 10 mM, and aliquoting and storing it at −20°C. Aliquots were diluted in physiological saline to a final concentration of 5 µM before use in experiments.

### Electrophysiology

The STNS preparation was superfused continuously with chilled *C. borealis* physiological saline (8 − 10°C), or Gly^1^-SIFamide (5 µM) diluted in saline. A switching manifold enabled uninterrupted superfusion of the STNS preparation during solution changes. Extracellular activity was recorded from nerves with custom stainless-steel pin electrodes and a model 1700 A-M Systems Amplifier. The stomatogastric ganglion (STG) was desheathed to gain access for intracellular recording of STG somata. Light transmitted through a dark field condenser (MBL-12010 Nikon Instruments) provided visualization of STG neuron somata. For intracellular recordings of STG neurons, sharp-tip glass microelectrodes pulled using a glass electrode puller (P-97, Flaming/Brown microelectrode puller, Sutter Instrument Co.) were used. Microelectrodes were filled with a squid internal solution (see Solutions) (resistance: 20 – 40 MΩ) (Hooper et al., 2015). Axoclamp 900A amplifiers (Molecular Devices) were used to collect intracellular recordings in current clamp mode. Recordings were collected before, during, and after the transection of both inferior and superior oesophageal nerves (*ion* and *son,* respectively), to isolate the STG from descending neuromodulatory inputs. Data presented were collected after transections were performed, unless otherwise noted. For intracellular recordings, STG neuronal cell bodies were identified using extracellular nerve recordings and/or their interactions with other STG neurons. All recordings were collected using acquisition hardware (Micro1401; Cambridge Electronic Design) and software (Spike2 Version 8.02e; ∼5kHz sampling rate; Cambridge Electronic Design) and a laboratory computer (Dell).

A data set from Fahoum and Blitz (2021) was re-used in this study for different measurements than the previous study. As described, activity in the inferior cardiac (IC), lateral gastric (LG), or dorsal gastric (DG) neurons, or all three neurons simultaneously, was eliminated via hyperpolarizing current injection (−2 to −4 nA). Neurons were hyperpolarized sufficiently to eliminate spike-mediated and graded transmitter release (Fahoum and Blitz, 2021).

For the established biological model of MCN5 actions, it is necessary to eliminate LP activity to mimic the MCN1 glutamatergic inhibition of LP (Fahoum and Blitz, 2021). Therefore, for all experiments using bath-application of Gly^1^-SIFamide (5 µM), the lateral pyloric (LP) neuron was either hyperpolarized (−2 to −4 nA) or photoinactivated (Fahoum and Blitz, 2021; Snyder and Blitz, 2022). To selectively eliminate the influence of the single LP neuron or the two copies of the LPG neurons, neurons were impaled with a microelectrode that was tip-filled with AlexaFluor-568 hydrazide (10 mM in 200 mM KCl; Thermo Fisher Scientific) and backfilled with squid internal solution. The neurons were injected with −5 nA hyperpolarizing current (for either ∼ 30 min, or for 5 to 10 min and the dye then allowed to diffuse for an additional ∼30 min) to fill their soma and neurites with the AlexaFluor-568 dye. The STG was then illuminated with a Texas red filter set (560 ± 40 nm wavelength; Leica Microsystems, 3-7 min). The neurons were considered completely photoinactivated when the membrane potential reached 0 mV, and their action potentials were absent in either the lateral ventricular nerve (*lvn;* for LP) or the lateral posterior gastric nerve (*lpgn;* for LPG).

### Data analysis

#### Neuronal Activity Quantification

The number of bursts, burst duration (s), cycle period (s; start of one burst to the start of the subsequent burst of a reference neuron), number of spikes per burst, and firing frequency (Hz; [number of spikes per burst – 1]/burst duration) were quantified using a custom Spike2 script. Each activity parameter was averaged across 20 min, unless otherwise noted. All analyses were performed after the effects of Gly^1^-SIFamide application reached steady state. Steady state was noted when LPG elicited consistent dual-network bursting (e.g., Fig. 1Ciii) with consistent gastric mill-timed bursting of LG, IC, and DG (∼ 10 min from the start of Gly^1^-SIFamide bath application). The pyloric cycle period was quantified as the duration(s) from the start of one pyloric dilator (PD) neuron burst to the start of the subsequent PD burst (Bartos et al., 1999). In the Saline:*ions*/*sons* cut condition, in which the *ions/sons* were transected, the pyloric cycle period did not always express a full triphasic pattern, due to the lack of modulatory input (Hamood et al., 2015; Spencer and Blitz, 2016; Zhang et al., 2009). However, when this occurred, the pyloric rhythm returned during the Gly^1^-SIFamide bath application. A Saline:*ions/sons* cut condition was excluded from analysis if PD neuron bursting was intermittently active, was completely absent, or if the PD neurons were rhythmically active but had ≤ 2 action potentials per burst. Pyloric cycle periods were measured across the final 20 min of 30 min Gly^1^-SIFamide (5 µM) bath applications, and across 200 s windows for Saline:Intact STNS and the Saline:*ion/son* cut conditions.

Gastric mill-timed IC bursts were identified as those that had burst durations greater than 0.45 s (Blitz et al., 2019; Fahoum and Blitz, 2024). Multiple IC bursts with greater than 0.45 s burst durations, were grouped into a single “gastric mill” burst if the distance between each burst was less than 2 s, otherwise they were considered as individual bursts. If there was an IC burst with a burst duration less than 0.45 s between two gastric mill IC bursts that were less than 2 s apart, they were grouped together as one gastric mill burst. Within an IC gastric mill-timed burst, pyloric-timed IC bursts were identified if there was an interruption timed to a PD burst (e.g., Fig. 6Ai-ii). We identified LPG gastric mill-timed bursts (LPG slow bursts) as previously described (Fahoum and Blitz, 2021; Fahoum and Blitz, 2024; Snyder and Blitz, 2022). Briefly, from a histogram of all LPG inter-spike intervals (ISIs) made in Excel (Microsoft), peaks of intraburst (interval between spikes during a burst) and interburst (interval between spikes between burst) intervals were identified. The mean ISI was calculated between these two peaks and used to identify LPG bursts. A custom written Spike2 script was then used to identify LPG gastric mill-timed slow bursts and exclude pyloric-timed LPG bursts using PD activity as a reference (Fahoum and Blitz, 2021). Coefficient of variation (CV) of pyloric cycle periods was calculated as Standard Deviation / Mean.

### Analysis of pyloric cycle period relative to gastric mill neuron activity

To analyze pyloric cycle periods based on which gastric mill neurons were active, burst timings for each neuron over a 20 min window, unless otherwise indicated, were exported from Spike2 to MATLAB (MathWorks) for further analysis. A custom written MATLAB script categorized PD cycle periods according to which gastric mill neuron(s) bursts overlapped with each pyloric cycle. This included any amount of overlap between a pyloric cycle period and a gastric mill neuron burst. Pyloric cycle periods that did not overlap with any gastric mill neurons were placed into the SIFbaseline category, i.e., no gastric mill neurons active. Pyloric cycle periods assigned to a particular category were averaged for each preparation, and a total average was then calculated across all experiments for each category. In some preparations, no pyloric cycle periods were assigned to some of the categories. Specifically, for IC, LPG:IC, and LPG:DG:IC there were 9 to 10 preparations that had no cycle periods assigned to each of these categories while for the remaining 13 categories there were 0 to 3 preparations that had no pyloric cycles assigned to these categories. This data set (n=16/19 from Fahoum and Blitz, 2021) and the LPG:Intact and LPG:Kill condition experiments (n=9/9 from Fahoum and Blitz, 2024) were reanalyzed from previous studies. Two experiments from the original data set of 11 were excluded from analysis due to a lack of biphasic regulation in the LPG:Intact condition.

Due to the inter-preparation variability of the pyloric cycle period, to test for changes in regulation of the pyloric cycle period between LPG:Kill and LPG:Intact conditions, all pyloric cycle periods were normalized to the average pyloric cycle period in the SIFbaseline category in each experiment. All pyloric cycle periods after normalization were plotted as a histogram with a bin size of 3 ms for both conditions. Further, bin counts for each histogram were normalized to the peak count in that condition to compare the overall distribution of the two histograms. Normalization and histogram plotting were performed using a custom written MATLAB script.

### Figure preparation and statistical analysis

Burst and spike identification from raw data was performed with custom-written scripts in Spike2, and exported to either Microsoft Excel or MATLAB for further analysis. Pyloric cycle period histograms were plotted using a script written in MATLAB. Figures were created using graphs and recordings exported from MATLAB, SigmaPlot (Systat, v13), and Spike2 into CorelDraw (Corel, v24). Statistical analyses were performed using SigmaPlot software and R statistical software (v4.1.1). Data were analyzed for normality using the Shapiro-Wilk test to determine whether a non-parametric or parametric test would be used. Paired *t* test, One Way Repeated Measures ANOVA, mixed model ANOVA, and *post hoc tests* were used as noted. For all tests, we used an alpha level of 0.05. Data are presented as mean ± SE.

### Code accessibility

Spike2 and MATLAB scripts used for analysis are available at https://github.com/blitzdm/GnanabharathiFahoumBlitz2024.

## Results

### Gly^1^-SIFamide elicits a variable pyloric rhythm

The modulatory projection neuron MCN5 elicits a unique gastric mill rhythm that can be reproduced in vitro by bath application of the MCN5 neuropeptide Gly^1^-SIFamide plus elimination of LP neuron activity (Blitz et al., 2019; Fahoum and Blitz, 2021; Fahoum and Blitz, 2024). MCN5 inhibits LP via its cotransmitter glutamate, but Gly^1^-SIFamide bath application excites LP, likely mimicking the actions of a second Gly^1^-SIFamide-containing input to the STG (Blitz et al., 2019; Fahoum and Blitz, 2021). Therefore, we hyperpolarized or photoinactivated the LP neuron to enable LPG dual-network activity and better mimic the MCN5-elicited motor pattern. In *C. borealis*, LPG is typically only active with the pyloric network due to its electrical coupling with the rest of the pyloric pacemaker ensemble (Fig. 1B-C) (Shruti et al., 2014), but switches into dual-network activity characterized by longer-duration, slower gastric mill-timed bursts alternating with shorter duration, faster pyloric-timed bursts during Gly^1^-SIFamide application (Fig. 1Ciii) (Blitz et al., 2019; Fahoum and Blitz, 2021; Fahoum and Blitz, 2024). The unique Gly^1^-SIFamide gastric mill rhythm is characterized by co-activity of LG and IC neurons that are primarily out of phase with DG neuron activity, followed by LPG neuron gastric mill activity (Blitz et al., 2019; Fahoum and Blitz, 2024) (Fig. 1C). Anecdotally, the pyloric rhythm cycle period varies throughout the Gly^1^-SIFamide gastric mill rhythm, with a correlation between IC burst duration and extended duration pyloric cycles (Blitz et al., 2019; Fahoum and Blitz, 2021). Here we investigate in detail the timing relationship between pyloric and gastric mill rhythms in the Gly^1^-SIFamide modulatory state.

**Figure 1.**
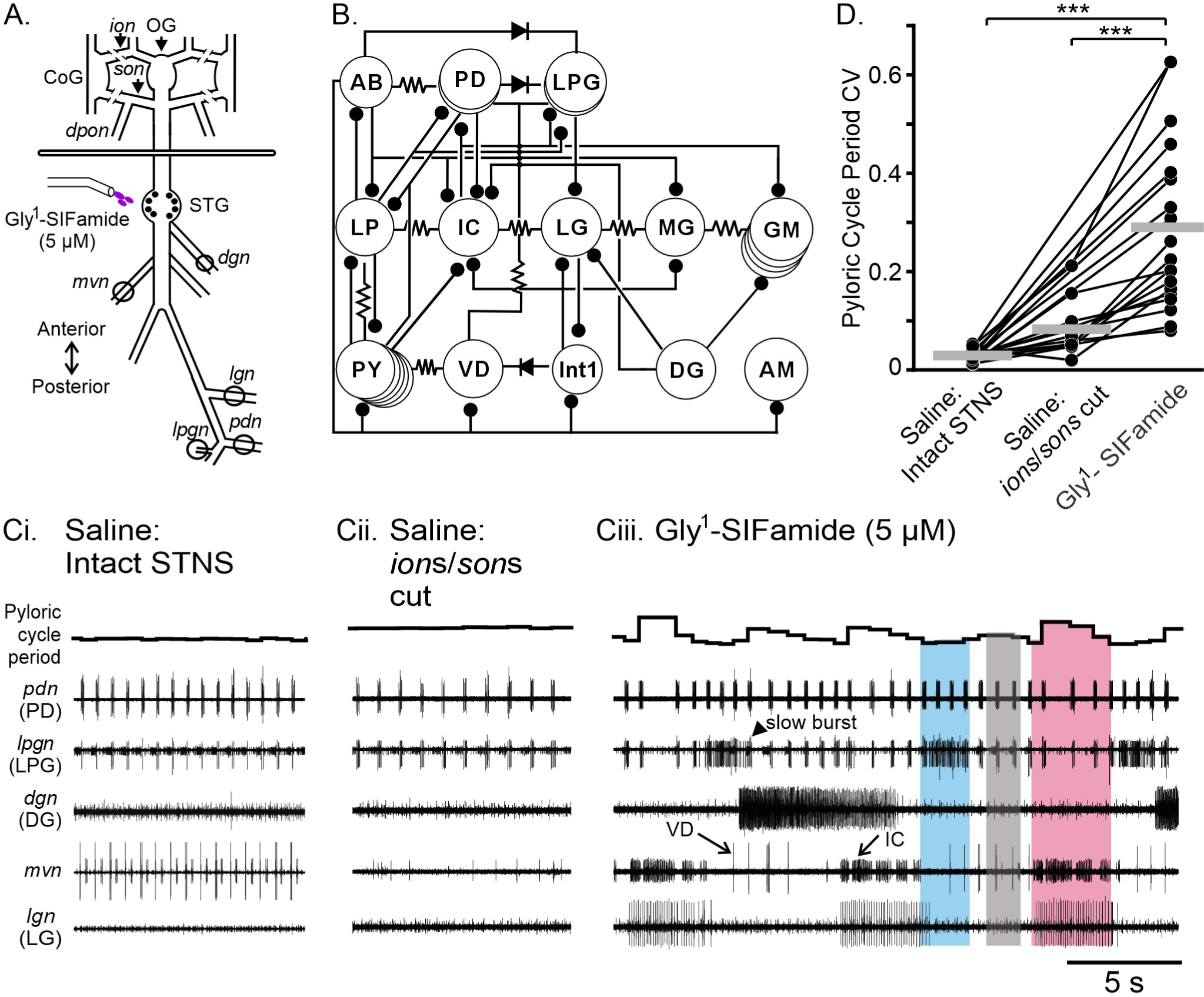
Gly^1^-SIFamide elicited a variable pyloric rhythm. ***A*,** Schematic of the isolated stomatogastric nervous system (STNS) of the crab, *Cancer borealis*. The STNS consists of the paired commissural ganglia (CoG), the single oesophageal ganglion (OG) and the single stomatogastric ganglion (STG), plus motor and connecting nerves. The STG consists of neuropil and the cell bodies of both the gastric mill and pyloric network neurons. The line breaks on the *ion* and *son* indicate where they were cut to isolate the STG network (Saline:*ions/sons* cut). Two compartments were created with a Vaseline wall across the dish and separately superfused. Gly^1^-SIFamide (5 µM) was bath-applied selectively to the posterior compartment containing the STG. ***B*,** Schematic of the pyloric and gastric mill connectome. Resistor symbols indicate electrical coupling, diode symbols indicate rectification of electrical coupling, and ball and stick symbols indicate chemical inhibition. Colors indicate neurons that participate in the Gly^1^-SIFamide rhythm and are discussed in this study. ***C*,** Representative electrophysiological traces of the pyloric (*pdn*, *mvn, lpgn*) and gastric mill (*mvn, dgn* and *lgn*) networks in an example experiment in Saline:Intact STNS (**Ci**), Saline:*ions/sons* cut (**Cii**) and bath-applied Gly^1^-SIFamide (5 µM) (**Ciii**) conditions. The instantaneous pyloric cycle period is plotted at the top of each set of traces. All conditions are from the same experiment. The colored boxes indicate three levels of pyloric cycle period, overlapping with three distinct phases of the Gly^1^-SIFamide gastric mill rhythm (blue: LPG activity; pink: IC/LG activity; grey: baseline Gly^1^-SIFamide modulation when no gastric mill neurons are active; see also Fig. 2). ***D*,** The coefficient of variation (CV) of the pyloric cycle period is plotted for three experimental conditions: 1) in saline prior to isolation of the STG network (Saline:Intact STNS, n = 19), 2) in saline with *ion*s*/son*s cut to isolate the STG network from descending modulatory inputs (Saline:*ion*s/*son*s cut, n = 12) and 3) during 5 µM Gly^1^-SIFamide bath application (SIFamide, n = 19). Each dot represents individual experiments and lines connecting across the experimental conditions depict data points from the same preparation. The different n-value for *ion*s/*son*s cut is due to the lack of a pyloric rhythm in seven preparations in this condition. In the absence of modulatory inputs, the pyloric rhythm sometimes shuts off (Zhang et al., 2009; Hamood et al., 2015). ***p < 0.001, One Way RM ANOVA, Holm-sidak *post hoc*. Neurons: AB, anterior burster; AM, anterior median; DG, dorsal gastric; GM, gastric mill; IC, inferior cardiac; Int1, Interneuron 1; LG, lateral gastric; LP, lateral pyloric; LPG, lateral posterior gastric; MG, medial gastric; PD, pyloric dilator; PY, pyloric; VD, ventricular dilator. Nerves: *dgn*, dorsal gastric nerve; *ion*, inferior oesophageal nerve; *lgn*, lateral gastric nerve; *lpgn*, lateral posterior gastric nerve; *lvn*, lateral ventricular nerve; *mvn*, median ventricular nerve; *pdn*, pyloric dilator nerve; *son*, superior oesophageal nerve; *stn*, stomatogastric nerve.

We first quantified the variability of the pyloric rhythm by measuring the coefficient of variation (CV) in control and during the Gly^1^-SIFamide gastric mill rhythm. To eliminate confounds due to a potentially variable set of endogenously active modulatory inputs, we transected the *ion*s and *son*s to obtain the same baseline condition across preparations. However, this often results in a very slow pyloric rhythm, or the pyloric rhythm shutting off entirely (Hamood et al., 2015; Zhang et al., 2009). Therefore, to compare the variability of the pyloric rhythm during Gly^1^-SIFamide to a similarly robust pyloric rhythm, we included a comparison of the pyloric rhythm in saline in the intact STNS, in addition to during saline superfusion after *ion*/*son* transection. In an example experiment, the pyloric cycle period was approximately 1.6 s, with a CV of 0.02 (Fig. 1Ci, PD activity in the *pdn,* instantaneous cycle period plotted above traces). After *ion*/*son* transection, the pyloric cycle period increased to ∼ 2.4 s and a CV of 0.05 (Fig.1 Cii). During Gly^1^-SIFamide, a gastric mill rhythm was activated, evident in LG and DG neuron bursting (Fig. 1Ciii, *lgn, dgn* recordings) and longer duration gastric mill-timed bursting in the IC neuron (*mvn*). Additionally, the LPG neuron switched from pyloric-only activity (Fig. 1Ci-ii; *lpgn*, Saline conditions) to dual pyloric and gastric mill (slow) bursting (Fig. 1Ciii). Although the average pyloric cycle period in Gly^1^-SIFamide (1.6 s) was similar to the intact STNS in this example, there was a visually evident increase in variability (CV = 0.22) (Fig. 1Ciii). The increased variability appeared to be due to three levels of pyloric cycle period (Fig. 1Ciii, colored boxes, discussed in more detail below). When quantified across preparations, we found a consistent increase in pyloric cycle period variability in Gly^1^-SIFamide compared to Saline, including both Saline:Intact STNS and after Saline:*ion*/*son* cut (Fig. 1D) (One Way RM ANOVA, F(18,2) = 30.09, p < 0.001, Table 1). In some experiments there was no active pyloric rhythm to quantify in the Saline:*ions/sons* cut condition and thus points between the Saline:Intact STNS and Gly^1^-SIFamide conditions are connected without an intermediate (Saline:*ion*s/*son*s cut) datapoint (see Methods). Thus, compared to the pyloric rhythm when influenced by endogenously active modulatory inputs (Saline:Intact STNS), or a slower pyloric rhythm in the absence of modulatory inputs, the pyloric rhythm cycle period was more variable in the Gly^1^-SIFamide modulatory state.

**Table 1.** Pyloric cycle period CV in saline (intact STNS or with *ion*s/*son*s transected) and during bath application of Gly^1^-SIFamide (5 μM).

| Pyloric Cycle Period CV<br>Mean ± SEM |  |  |  | p-value |  |  |
| --- | --- | --- | --- | --- | --- | --- |
| Saline: Intact<br>STNS<br>(n = 19) | Saline: *ion/son*<br>cut<br>(n = 12) | Gly<sup>1</sup>-SIFamide<br>(n = 19) | F<br>(df) | Saline: Intact<br>STNS vs<br>SIFamide | Saline: *ion/son*<br>cut vs<br>SIFamide | Saline: Intact<br>STNS vs<br>Saline: *ion/son*<br>cut |
| 0.03 ± 0.002 | 0.08 ± 0.01 | 0.29 ± 0.04 | 30.09<br>(18,2) | <0.001 | <0.001 | 0.182 |

One-way repeated measures ANOVA, Holm-Sidak *post hoc*; CV, coefficient of variation; df, degrees of freedom

### Biphasic Regulation of the Pyloric Rhythm

The larger pyloric cycle period CV during Gly^1^-SIFamide application appeared to be due to rhythmic changes in cycle period coinciding with activity in some of the gastric mill neurons. Specifically, in example experiments, the pyloric cycle period was longer during times when the IC and LG neurons were active (Figs. 1Ciii, 2A, pink box) relative to pyloric activity when no gastric mill neuron was active (Figs. 1Ciii, 2A; SIFbaseline; grey box; dotted horizontal line across plot of instantaneous cycle period). Shorter duration pyloric cycle periods relative to SIFbaseline coincided with LPG neuron slow bursts (Figs. 1Ciii, 2A; blue box).

**Figure 2.**
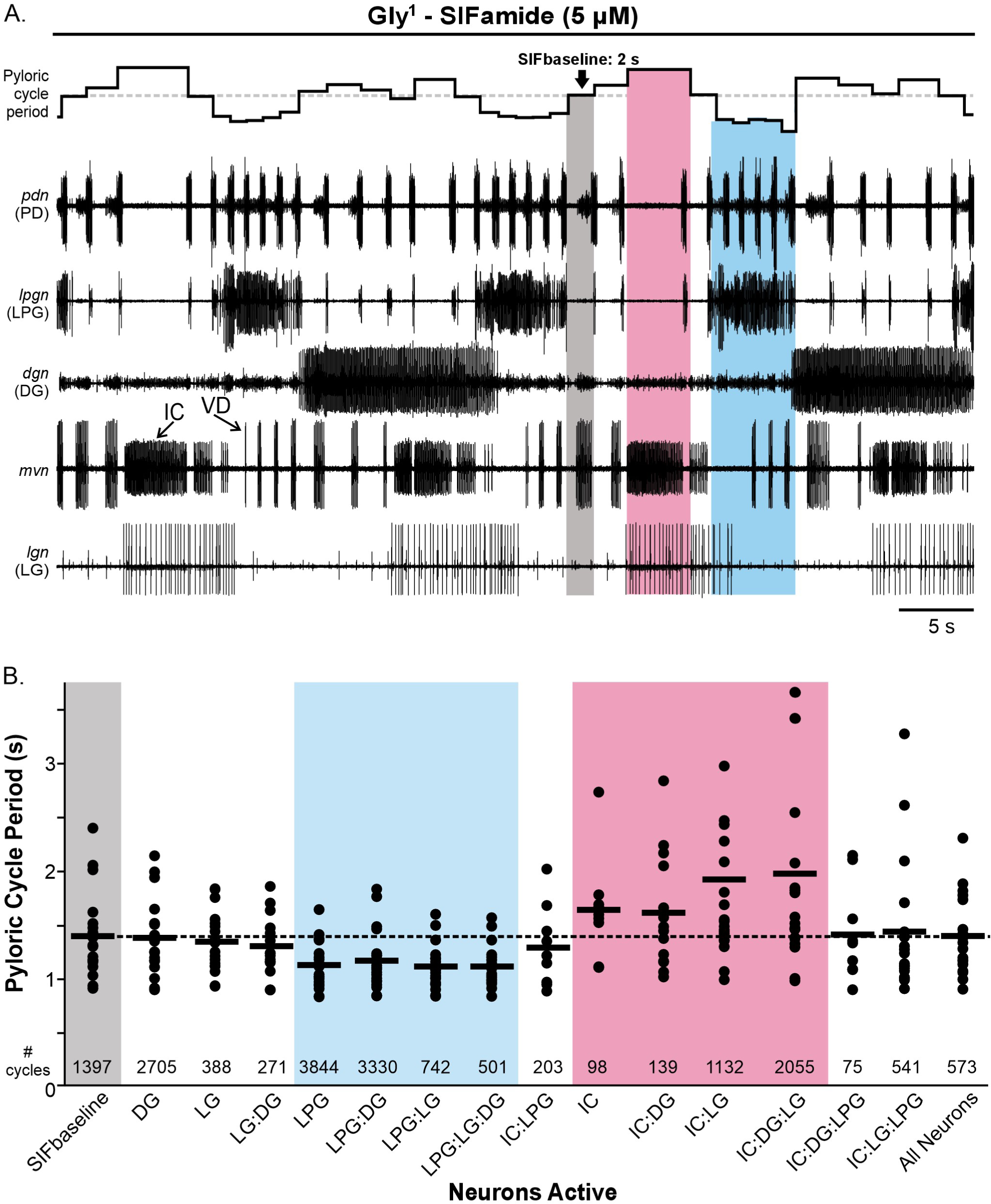
Pyloric cycle periods occurred at three different levels during the Gly^1^-SIFamide gastric mill rhythm. ***A*,** The pyloric rhythm occurs with a shorter cycle period during LPG activity (blue box), with a longer cycle period during LG:IC activity (pink box) and an intermediate level in the absence of any gastric mill-timed bursting (SIFbaseline; grey box). Representative traces show an example rhythm during Gly^1^-SIFamide bath application, with instantaneous pyloric cycle period plotted at the top. Dotted line across instantaneous cycle period plot indicates SIFbaseline cycle period. ***B*,** The average pyloric cycle period is plotted for no gastric mill neuron activity (SIFbaseline; grey box) and during activity of each combination of 1, 2, 3, and 4 gastric mill neurons (n = 19). Each dot represents the average pyloric cycle period during that particular neuron activity across a 20 min analysis window during Gly^1^-SIFamide steady state in a single experiment. Black bars indicate the mean pyloric cycle period across all experiments during that neuronal activity. Numbers at the bottom of the graph depict the total number of pyloric cycles contributing to the averages for that neuronal activity. Dotted line represents average pyloric cycle period during SIFbaseline (grey box). Blue box highlights neuronal activity combinations that include LPG, and pink box highlights combinations that include IC.

To further examine the relationship between pyloric cycle period and gastric mill neurons, we measured all pyloric cycle periods across the final 20 min steady-state of 30 min Gly^1^-SIFamide (5 µM) bath applications and grouped them by the gastric mill neurons that were active. Considering the four gastric mill neurons (LG, LPG, IC, DG), plus the SIFbaseline condition in which no gastric mill neurons are active, there are 16 possible combinations of active gastric mill neurons during which a pyloric cycle period might have occurred (Fig. 2B). We focused only on gastric mill-timed bursts (see Methods; Fahoum and Blitz 2024), including for IC and LPG which express dual-network activity. The average cycle period occurring during each neuron or combination of neurons active, per experiment (black dots) and across all experiments (black bars) are plotted for the 16 different combinations (Fig. 2B). Across 19 preparations, there were some preparations in which no pyloric cycle periods overlapped with a combination of neurons. Specifically in the IC, LPG:IC, and LPG:DG:IC categories there were between 9 and 10 preparations with no pyloric cycle periods falling into these categories. In the 13 remaining categories there were 0 to 3 preparations that had no pyloric cycles in these categories. The total number of pyloric cycles that occurred during each condition, across 19 preparations, are indicated above the x axis labels (Fig. 2B; # cycles).

We first visually assessed cycle periods as a function of gastric mill neurons active, using a dashed line to highlight the average cycle period across preparations during the SIFbaseline, when no gastric mill neurons were active (grey region, dashed line). Relative to this baseline, the average cycle period per experiment (dots) tended to be shorter in duration than SIFbaseline during LPG activity, whether it was LPG alone or in combination with DG and/or LG neurons (blue region), including the cumulative average cycle period across preparations (black bars) (n = 16-19; Fig. 2B). Conversely, average cycle periods during combinations of IC, DG, and LG activity (pink region) tended to be longer, and thus above SIFbaseline in the graph, including the cumulative average cycle period (IC, n = 9; IC:DG, IC:LG, IC:DG:LG, n = 19). This aligns with an earlier study in which longer pyloric cycle periods correlated with longer duration IC bursts (Blitz et al., 2019). Based on the number of cycles, IC gastric mill-timed bursts most often coincided with LG activity (Fig. 2B, # cycles). Overall, this representation of cycle period as a function of gastric mill neuron activity suggests that LPG, IC, and LG activity may be responsible for the increased variability in pyloric cycle period during the Gly^1^-SIFamide gastric mill rhythm compared to saline (Fig. 1D). To analyze this data set statistically, we used a mixed model ANOVA to accommodate the large number of conditions, i.e., combinations of gastric mill neurons active, plus missing values for some combinations (n = 9-19).

For the statistical analysis of gastric mill neuron activity on the pyloric cycle period, we considered each gastric mill neuron to be a factor with a full 4-factor within-subjects mixed model ANOVA, including all higher-order interactions. This model indicated statistically significant effects of gastric mill neuron combinations on pyloric cycle period variability (Χ^2^_df = 15_ = 102.57, p < 0.0001). The model complexity was then iteratively paired down, eliminating non-significant higher-order interactions (Table 2). The most parsimonious model that still explained the variability in pyloric cycle period included single factors LPG, IC, LG and 2-factor terms IC:LPG and IC:LG (Table 2). In this model, there were significant interactions between LG and IC (Χ^2^_df = 239_ = 3.07, p = 0.002) and between LPG and IC (Χ^2^_df = 238_ = −2.38, p = 0.02). Specifically, LG had an effect on pyloric cycle period when IC was firing a gastric mill burst (p = 0.0011, Table 2) but not without IC gastric mill activity (p = 0.40, Table 2). However, IC had an effect on cycle period with (p<0.0001) or without LG activity (p=0.48, Table 2), although a weaker effect in the absence of LG. This points to IC playing an important role in the longer cycle pyloric cycle periods, with possible assistance from LG. For the LPG and IC interaction, LPG had an effect on pyloric cycle period with (p < 0.0001) or without (p = 0.0004, Table 2) IC generating a gastric mill burst. Similarly, IC affected the pyloric cycle period with (p = 0.008) or without (p < 0.0001, Table 2) LPG firing a gastric mill burst. Thus, from the overlap in timing and the preceding statistical analysis, it appears that IC (possibly LG) and LPG are responsible for the increased and decreased cycle period, respectively, and the consequent higher cycle period variability during the Gly^1^-SIFamide gastric mill rhythm. We therefore tested the roles of IC, LG, and LPG in regulating pyloric cycle period by manipulating their activity.

**Table 2.** Statistical analysis of active gastric mill neuron contributions to pyloric cycle period variability during Gly^1^-SIFamide (5 μM) application.

| Model Complexity |  | X<sup>2(a,b)</sup>, df | p-value |
| --- | --- | --- | --- |
|  |  | t<sup>c</sup>, df |  |
| 4-factor interaction term removed |  | 0.0612<sup>a</sup>, 1 | 0.8047 |
| All 3-factor interaction terms removed |  | 2.9444<sup>b</sup>, 4 | 0.5672 |
| <u>Final Model:</u> |  | 1.019<sup>b</sup>, 5 | 0.961 |
| Includes LPG, LG, IC, LPG:IC, LG:IC |  | 4.015<sup>a</sup>, 10 | 0.946 |
| Post hoc<br>comparisons<br>of significant<br>2-factor<br>interactions<br>in final<br>model | LG with IC Present | -3.30<sup>c</sup>, 240 | 0.011 |
|  | LG with IC Absent | 0.84<sup>c</sup>, 239 | 0.40 |
|  | IC with LG Present | -7.15<sup>c</sup>, 238 | <0.0001 |
|  | IC with LG Absent | -1.99<sup>c</sup>, 241 | 0.048 |
|  | LPG with IC Present | -6.35<sup>c</sup>, 238 | <0.0001 |
|  | LPG with IC Absent | -3.57<sup>c</sup>, 238 | 0.0004 |
|  | IC with LPG Present | -2.69<sup>c</sup>, 239 | 0.008 |
|  | LPG with IC Absent | -6.23<sup>c</sup>, 238 | <0.0001 |

4-factor within-subjects mixed model ANOVA with iterative elimination of non-significant higher-order interactions. Models were compared to full model<sup>a</sup> or previous level of complexity<sup>b</sup> using a likelihood ratio test. <sup>c</sup>post hoc comparisons of estimated marginal means.

### IC, but not LG, is responsible for longer pyloric cycle periods

We first addressed whether IC and/or LG were necessary for longer pyloric cycle periods during the Gly^1^-SIFamide gastric mill rhythm. The pyloric cycle period was variable in control conditions as is evident in two example experiments (Fig. 3Ai, Bi, pdn, instantaneous cycle period plot above traces). This included longer pyloric cycle periods during gastric mill-timed coactivity in LG and IC (Fig. 3; pink boxes). Identified IC gastric mill bursts (see Methods) (Fahoum and Blitz, 2024), are marked with brackets. In one instance in the time range shown, when LG fired a gastric mill burst, but IC produced only pyloric-timed activity that did not reach the level of a gastric mill-timed burst (Fig. 3Ai; grey box), there was no extension of the pyloric cycle period. This is consistent with the statistical analysis of data from figure 2B, that LG had an effect on pyloric cycle period, only when IC was also active (in gastric mill time). To test whether IC and/or LG was responsible for the longer pyloric cycle periods, we reanalyzed a data set from Fahoum and Blitz (2021) in which hyperpolarizing current (−2 to −4 nA) was injected into either IC or LG neurons. In the example experiments, the much longer cycle periods were eliminated when either LG or IC was hyperpolarized (Fig. 3Aii, Bii). When IC was hyperpolarized, LG bursting persisted, and was qualitatively similar to pre-hyperpolarization (Fig. 3Ai-ii). However, when LG was hyperpolarized, IC did not generate any gastric mill-timed bursts, instead its activity was entirely pyloric-timed (Fig. 3Bii). Longer pyloric cycle periods, and therefore larger cycle period variability, returned post hyperpolarization, as did IC gastric mill-timed bursting upon removal of hyperpolarizing current into LG (Fig. 3Aiii, Biii).

**Figure 3.**
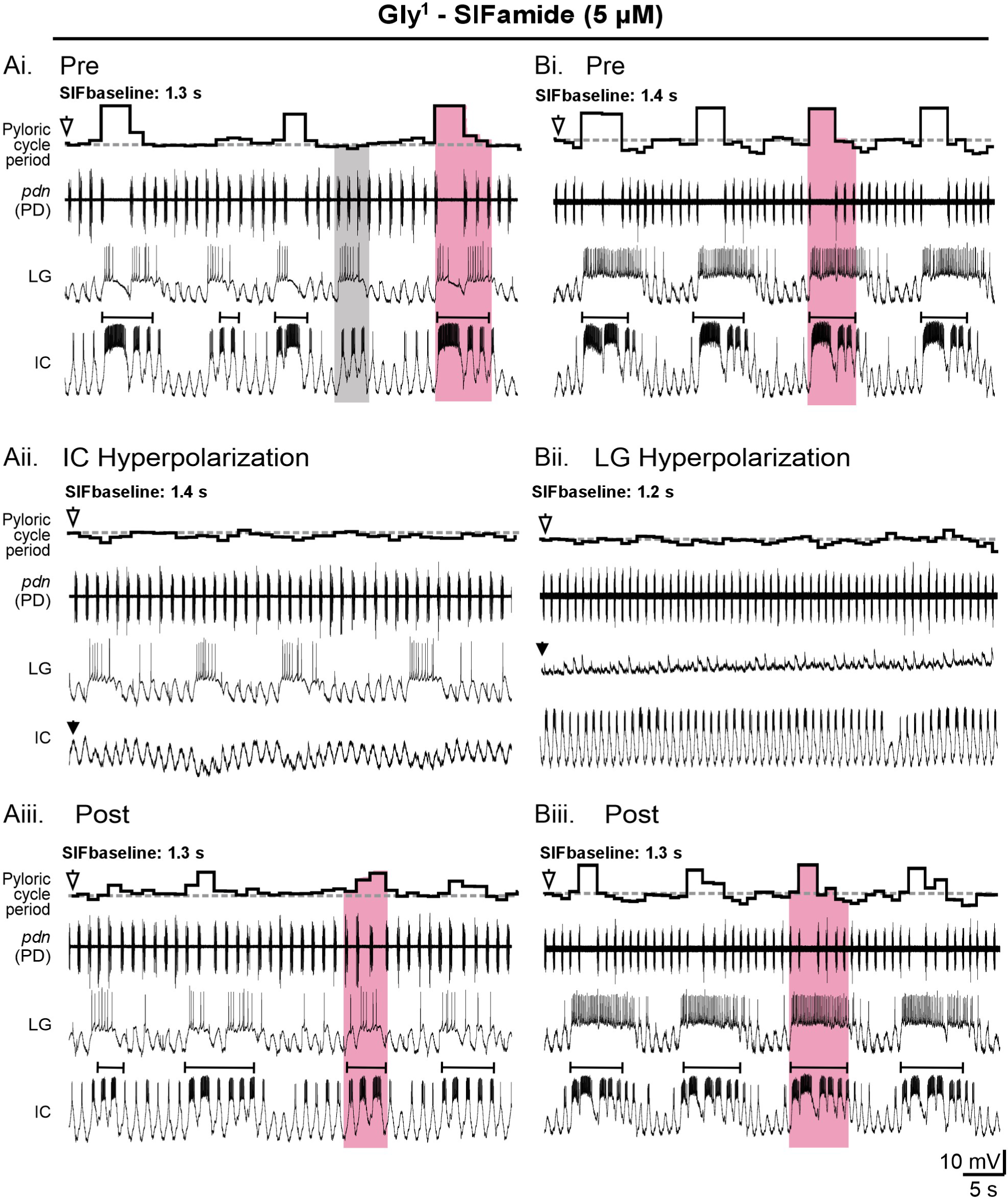
IC or LG hyperpolarization eliminated longer cycle periods and decreased pyloric cycle period variability. ***A***, Extracellular and intracellular recordings monitor the pyloric rhythm (*pdn*) and gastric mill-timed bursting in the IC and LG neurons in control (Pre; ***Ai***, ***Bi***; Top), during IC (***Aii***) or LG (***Bii***) hyperpolarization, and after hyperpolarization (Post; ***Aiii***, ***Biii***), all during Gly^1^-SIFamide bath application. A and B are from different experiments. Pyloric cycle period on top of the extracellular *pdn* recording plots the instantaneous pyloric cycle period during each condition. Note the absence of longer pyloric cycle periods during both LG and IC hyperpolarization (pink boxes indicate LG:IC activity, grey box indicates LG activity without an IC gastric mill burst (Ai). IC gastric mill bursts are identified by brackets above the IC recordings. Downward filled arrowheads indicate hyperpolarizing current injection (Aii and Bii). Downward white arrowheads above instantaneous pyloric cycle period indicate the SIFbaseline cycle period (dashed line).

To quantify the results of hyperpolarizing IC and LG, we measured pyloric cycle period CV before (Pre), during (Hype) and after (Post) hyperpolarization (Fig. 4A). The pyloric cycle period CV decreased when IC was hyperpolarized (Fig. 4Ai; black dots and connecting lines indicate results from individual experiments, grey bars indicate averages across experiments) (One Way RM ANOVA, F(5,2) = 7.43, p = 0.01, Table 3) and when LG was hyperpolarized (Fig. 4Aii; One Way RM ANOVA, F(3,2) = 6.09, p = 0.04, Table 3). We also determined that the pyloric cycle period CV was not altered when DG was hyperpolarized (Fig. 4Aiii; One Way RM ANOVA, F(3,2) = 1.18, p = 0.37, Table 3) but it was lower when all three gastric mill neurons (IC, LG and DG) were hyperpolarized (Fig. 4Aiv; One Way RM ANOVA, F(5,2) = 9.13, p = 0.01, Table 3). In the cumulative data, a lower variability when IC or LG or the three gastric mill neurons were hyperpolarized further indicated that IC and/or LG, but not DG, were responsible for regulating pyloric cycle period. When LG was hyperpolarized, IC gastric mill bursting was eliminated in the example experiment in figure 3B. Therefore, we aimed to quantify IC and LG activity across all experiments when either LG or IC was hyperpolarized. However, in 3/4 experiments there were no IC gastric mill bursts when LG was hyperpolarized. Therefore, instead of quantifying IC gastric mill bursts, we reported the number of IC gastric mill bursts before (Pre), during (Hype) and after (Post) LG hyperpolarization (Fig. 4B). There was a reversible decrease in IC gastric mill bursts with LG hyperpolarization (Fig. 4B) (One Way RM ANOVA, F(3,2) = 22.75, p = 0.002, Table 3). In contrast, when IC was hyperpolarized, LG activity, including burst duration (Fig. 4Ci), firing frequency (Fig. 4Cii) and number of spikes per burst (Fig. 4Ciii) did not change (One Way RM ANOVA; F(5,2) = 0.04 – 1.52; all conditions, p > 0.05, Table 3). Thus, LG impacts IC activity, but IC does not alter LG activity. It therefore appears likely that hyperpolarizing LG eliminated extended pyloric cycle periods via elimination of IC gastric mill bursting. In combination with a significant contribution of LG to pyloric cycle period variability, only when IC is active (above) these results indicate that IC, but not LG, gastric mill-timed bursts are responsible for the longer pyloric cycle periods.

**Figure 4.**
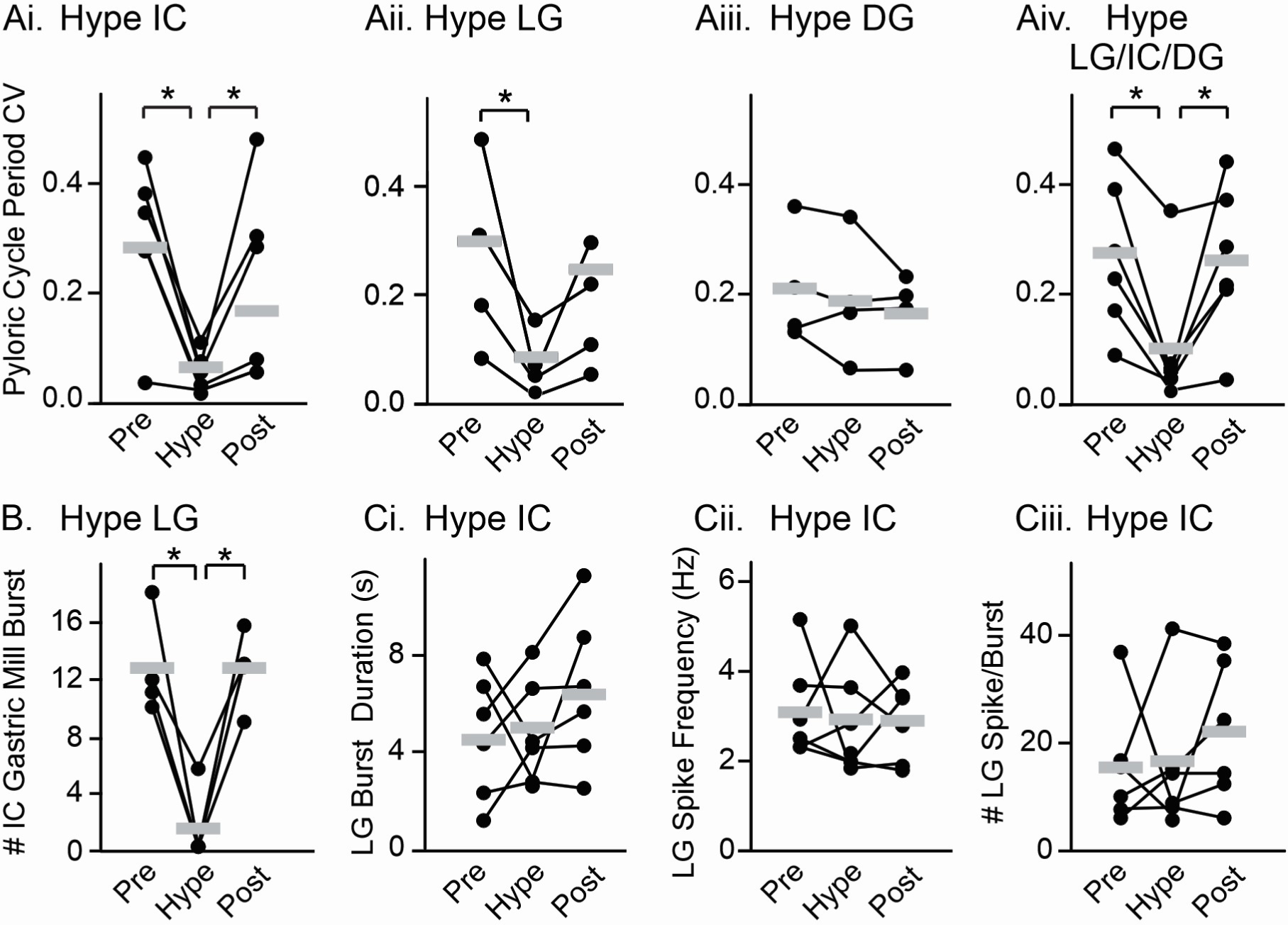
IC hyperpolarization decreased pyloric cycle period variability without altering LG activity. ***A***, The average CV of pyloric cycle period across 200 sec windows is plotted for IC (***Ai***, n = 6), LG (***Aii***, n = 4), DG (***Aiii***, n = 4) or all three neurons (***Aiv***, n = 6) before (Pre), during (Hype) and after (Post) hyperpolarization. ***B***, The number of IC gastric mill bursts (burst duration > 0.45 s) occurring before (Pre), during (Hype) and after (Post) LG hyperpolarization is plotted (n = 4). ***C***, LG burst duration (***Ci***), firing frequency (***Cii***) and number of spikes per burst (***Civ***) before (Pre), during (Hype) and after (Post) IC hyperpolarization are plotted (n = 6). For all graphs, data points represent individual experiments connected by lines across conditions. Grey bars indicate the average across experiments. *p < 0.05, One Way RM ANOVA, Holm-sidak *post hoc* test.

**Table 3.**
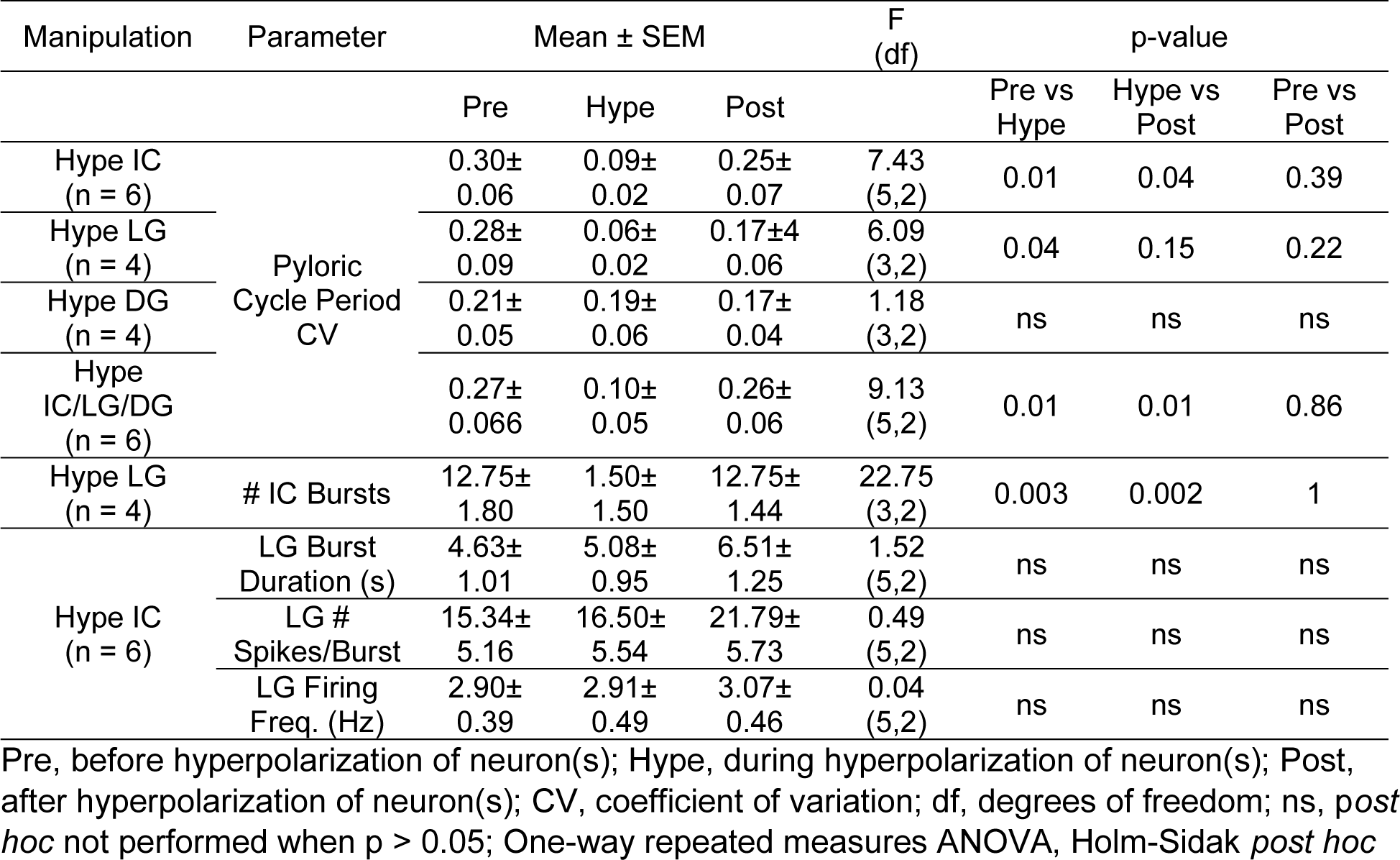
Pyloric cycle period CV and IC and LG gastric mill activity in Gly^1^-SIFamide (5 μM) before, during, and after hyperpolarization of gastric mill neurons.

### LPG contributes to pyloric cycle period variability in Gly^1^-SIFamide

LPG gastric mill-timed activity correlated with decreased pyloric cycle periods (Figs. 1, 2, 5). LPG is electrically coupled to the other pyloric pacemaker neurons (Fig. 1B) (Marder and Bucher, 2007; Shruti et al., 2014) and thus hyperpolarizing current injection into the two LPG neurons could alter the pyloric cycle period. Therefore, to test whether LPG was responsible for decreased pyloric cycle periods relative to SIFbaseline (e.g., Fig. 5Ai), the two LPG neurons were photoinactivated (see Methods; LPG:Kill) to selectively remove them from the network, and pyloric cycle periods during Gly^1^-SIFamide bath application were compared between LPG:Intact and LPG:Kill. Photoinactivation of two neurons does not alter the response to bath-applied Gly^1^-SIFamide, and responses to two consecutive bath applications of Gly^1^-SIFamide are not different (Fahoum and Blitz, 2024).

**Figure 5.**
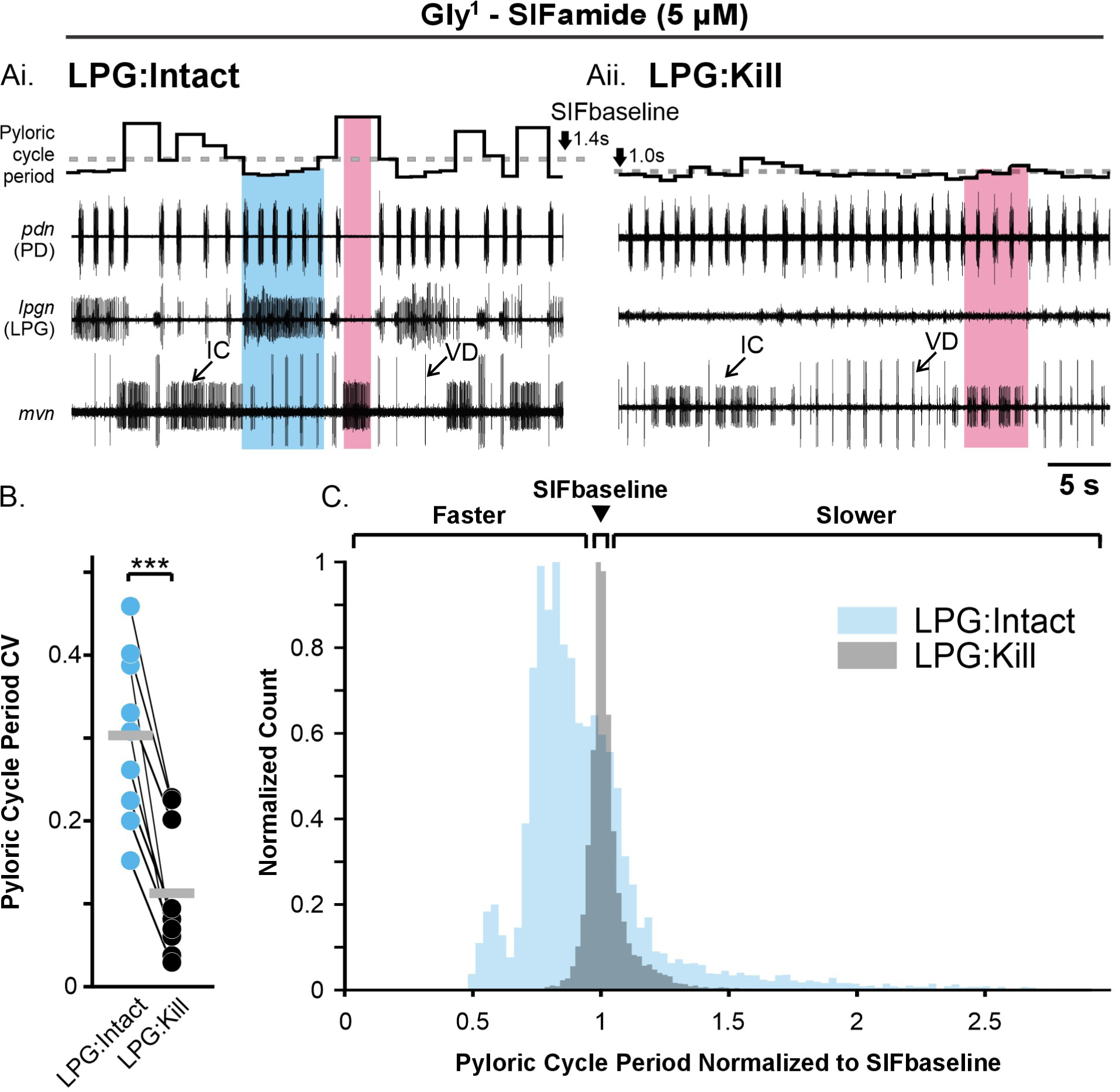
LPG photoinactivation decreased pyloric cycle period variability in Gly^1^-SIFamide. ***Ai,*** In control Gly^1^-SIFamide with LPG neurons intact, there are shorter cycle periods during LPG gastric mill-timed bursts (blue box), compared to SIFbaseline (dashed line) and longer cycle periods during IC activity (pink box). ***Aii***, After both LPG neurons were photoinactivated (see Methods), pyloric cycle period variability appeared lower. Pink box highlights an IC gastric mill burst (group of pyloric-timed bursts > 0.45 s duration). The small units in the *lpgn* recording are PY neurons. ***B***, The pyloric cycle period coefficient of variation (CV) is plotted for a 20 min window of steady state activity in the LPG:Intact (blue dots) and LPG:Kill (black dots) conditions (n = 9). Each pair of dots plus their connecting lines represents a single preparation. Grey bars indicate the average CV across experiments in each condition. ***p < 0.001, paired t-test. ***C***, Overlaid histograms (bin size: 3 ms) of all pyloric cycle periods across 20 min windows during steady state Gly^1^-SIFamide application in the LPG:Intact (blue) and LPG:Kill (grey) conditions (n = 9). Each individual cycle period was normalized to the average SIFbaseline of the same preparation. Bin count of each condition was normalized to the peak count of that condition to facilitate comparison of histograms. Brackets on top of histogram indicate shorter pyloric cycle periods, SIFbaseline (arrow) and longer pyloric cycle periods.

In the absence of the LPG neurons (LPG:Kill), the pyloric cycle period variability decreased (Fig. 5Aii) compared to the LPG:Intact condition. Eliminating LPG prevented the periodic decreases in pyloric cycle period. Across experiments, the pyloric cycle period CV was lower after LPG:Kill, compared to LPG:Intact (Fig. 5B; LPG:Intact [blue dots] CV = 0.30 ± 0.03; LPG:Kill [black dots] CV = 0.11 ± 0.03; n = 9; two-tailed paired t-test; t(8) = 7.78; p < 0.001). However, the decreased variability did not appear to be due to only elimination of shorter pyloric cycle periods. The longer duration pyloric cycle periods were also mostly eliminated in the LPG:Kill condition as can be seen in the example experiment (Fig. 5Aii). Assessing regulation of pyloric cycle period across preparations was complicated by the absence of LPG which eliminated several combinations of active gastric mill neurons. Thus, we chose instead to examine the distribution of cycle periods in the LPG:Intact vs LPG:Kill conditions. There is variability in SIFbaseline pyloric cycle period across preparations (e.g., Fig. 2B, grey box), and photoinactivation of the two LPG neurons shifted the SIFbaseline to shorter pyloric cycle periods (compare instantaneous pyloric cycle period plot in Fig. 5Ai-ii), likely due to removal of “electrical drag” on the pyloric pacemaker neuron AB (Kepler et al., 1990). Thus, to determine whether there was any change in pyloric cycle period regulation between the two conditions, we normalized each pyloric cycle period to the average SIFbaseline cycle period in that preparation and accumulated all normalized pyloric cycle period durations during each condition into their respective histograms (Fig. 5C) (LPG:Intact, blue and LPG:Kill, grey). We also normalized pyloric cycle period count per bin to the highest count for each condition to better enable comparison of the histograms. In the LPG:Intact condition, in addition to the peak around 1 (SIFbaseline, arrowhead), there were shorter pyloric cycle periods (<1; “Shorter” bracket) and a distributed set of bins at longer cycle periods (>1; “Longer” bracket) (Fig. 5C, blue histogram). However, in the LPG:Kill condition (Fig. 5C, grey histogram), there was a single grouping of bars centered around 1 (SIFbaseline) without a population of slower or faster cycle periods. Therefore, removing LPG from the network prevented the increase and the decrease in pyloric cycle period. Electrical coupling between LPG and the pyloric pacemaker can explain why the pyloric rhythm was faster during the strong depolarizations of LPG slow bursts. However, this does not explain why removing LPG also eliminated slowing of the pyloric rhythm. Further, we showed above that IC gastric mill bursting was responsible for extending the pyloric cycle period, slowing the rhythm (Figs. 3-4). This combination of findings led us to ask whether photoinactivation of LPG alters IC gastric mill activity.

### IC gastric mill bursts were not impacted by LPG photoinactivation

As discussed previously, IC displays both pyloric- and gastric mill-timed bursts, with a gastric mill burst often consisting of multiple pyloric-timed bursts (Fig. 6A). In an example expanded region, a single IC gastric mill burst (Fig. 6Ai, brackets above *mvn* recording) in the LPG:Intact condition consists of four pyloric-timed bursts (Fig. 6Ai, brackets below *mvn* recording). In the LPG:Kill condition, based on established criteria for IC gastric mill bursts (Blitz et al., 2019; Fahoum and Blitz, 2024), a single IC gastric mill burst is approximately the same duration as in the LPG:Intact condition, but consists of seven pyloric bursts, compared to four with LPG neurons intact (Fig. 6A). In a previous study, there was no difference in IC activity between LPG:Intact and LPG:Kill, including no difference in burst duration, firing frequency, and number of spikes per burst for gastric mill bursts and for IC pyloric bursts within IC gastric mill bursts (Fahoum and Blitz, 2024). Because our LPG:Intact, LPG:kill dataset was a subset from the previous study (n = 9/11, see Methods), we reanalyzed the data to verify the result in the specific experiments used here. We found no difference in number of IC gastric mill bursts (Fig. 6Bi; Table 4), IC gastric mill burst duration (s) (Fig. 6Bii; Table 4), number of IC spikes per gastric mill burst (Fig. 6Biii; Table 4), or firing frequency (Hz) (Fig. 6Biv; Table 4). There was also no difference in the number of IC pyloric bursts (Fig. 6Ci; Table 4), IC pyloric burst duration (s) (Fig. 6Cii; Table 4), or IC pyloric firing frequency (Hz) (Fig. 6Civ; Table 4), although there was a trend toward shorter pyloric-timed burst duration (Fig. 6Cii; Table 4), and there was a decrease in the number of spikes per pyloric-timed burst (Fig. 6Ciii; Table 4). The difference between these results and the previous study likely reflects the removal of two preparations in which there was little pyloric regulation in the control condition. For the nine preparations in which there was typical pyloric regulation in control, the tendency for IC pyloric bursts to be shorter, with fewer action potentials in the LPG:Kill condition fits with the loss of long-duration pyloric cycle periods. When longer pyloric cycle periods are eliminated, IC neuron pyloric bursts are shortened as they are interrupted by inhibition from the pyloric pacemaker ensemble at the start of each pyloric cycle (e.g., Fig. 6A). Importantly however, IC gastric mill bursts were not altered by LPG:Kill and IC firing frequency, a measure of the strength of neuronal activity, was not altered for IC gastric mill or pyloric bursts (Fig. 6Biv, Civ).

**Figure 6.**
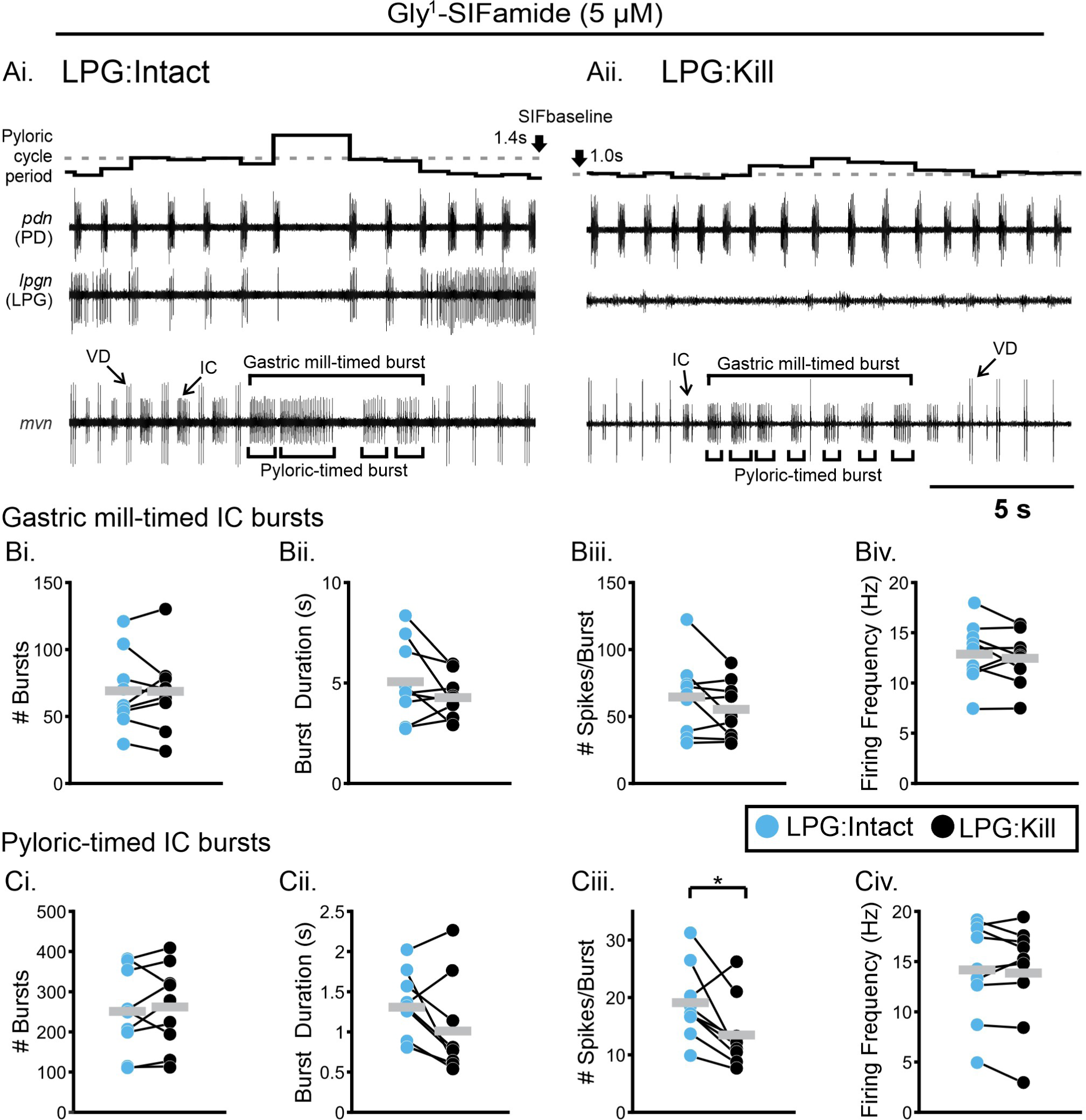
IC firing frequency was not affected by LPG photoinactivation. ***A***, Representative traces illustrate differences in IC gastric mill-timed activity (brackets on top of *mvn* recording) before (***Ai***, LPG:Intact) and after (***Aii***, LPG:Kill) photoinactivation of the two LPG neurons. Pyloric-timed IC bursts within each IC gastric mill burst were identified if there was a gap between IC action potentials that coincided with a PD neuron burst (brackets below *mvn* recording). ***B***, IC gastric mill-timed activity was quantified across 20 min windows during steady state Gly^1^-SIFamide bath application, including number of bursts (***Bi***), burst duration (***Bii***), number of spikes/burst (***Biii***), and firing frequency (***Biv***). ***C***, IC pyloric-timed activity within each gastric mill-timed burst was also quantified, including number of bursts (***Ci***) burst duration (***Cii***), number of spikes/burst (***Ciii***), and firing frequency (***Civ***). Blue dots indicate LPG:Intact condition and black dots indicate LPG:Kill condition. All dots represent the average within an experiment with lines connecting the two conditions in each experiment. Grey bars indicate averages across experiments. *p < 0.05, paired t-test.

**Table 4.**
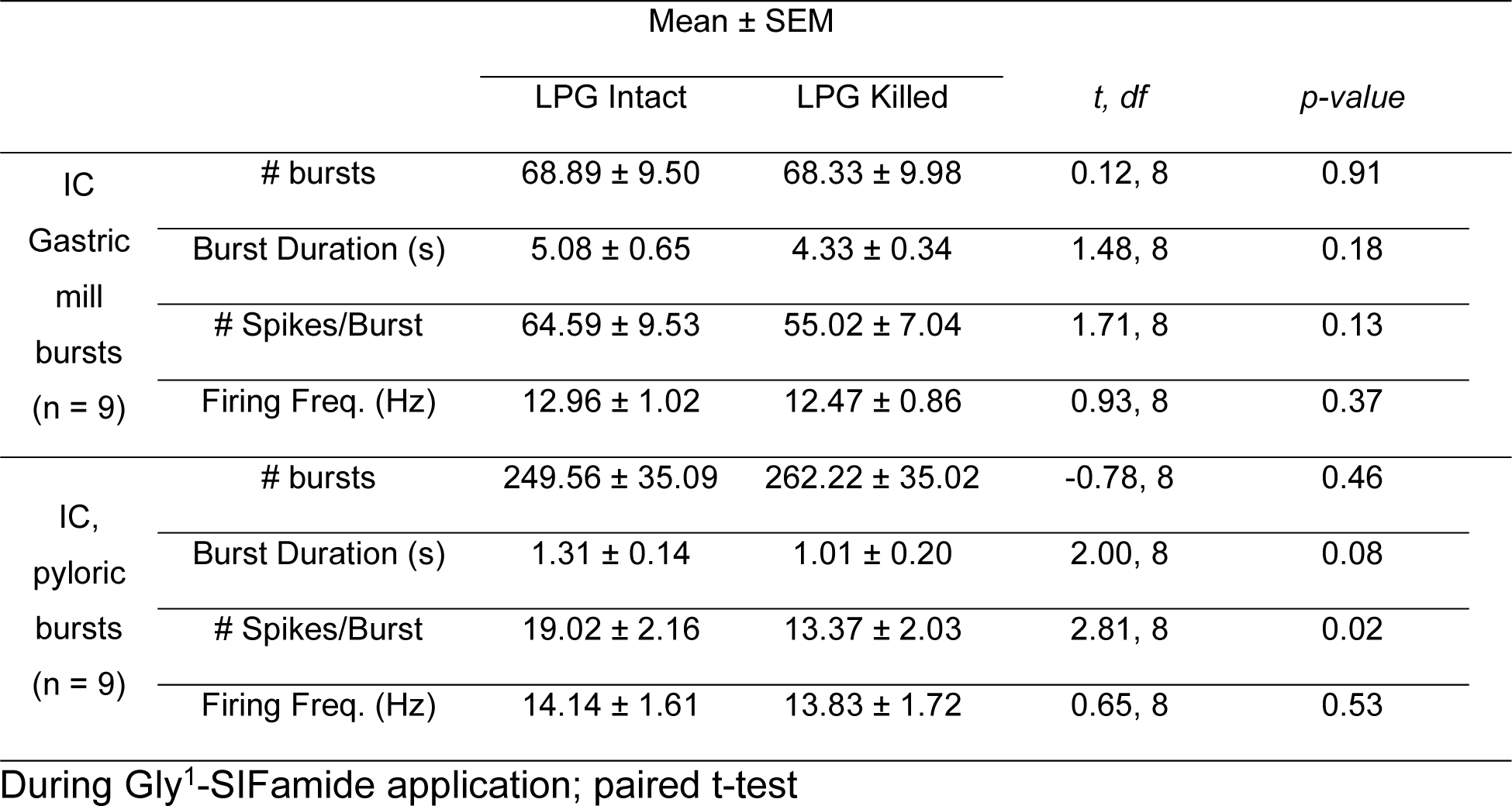
IC gastric mill- and pyloric-timed activity during LPG Intact and LPG photoinactivated (Killed) conditions in Gly^1^-SIFamide (5 µM).

### Same IC activity level does not slow the pyloric rhythm in the LPG:Kill condition

To determine the relationship between the strength (firing frequency) of IC gastric mill bursts and pyloric cycle period before versus after LPG:Kill, we plotted instantaneous pyloric cycle period against IC firing frequency (Hz) for all pyloric cycles across a 20 min steady state region of Gly^1^-SIFamide application for each of nine preparations (Fig. 7). Within a preparation, across the same IC firing frequencies, there were many longer duration pyloric cycle periods in LPG:Intact (blue circles) but many fewer longer duration cycles in the LPG:Kill condition (grey circles) (Fig. 7; n = 9). This indicated that LPG was playing a major role in the ability of IC to increase pyloric cycle period. The occasional ability of IC to increase pyloric cycle period in the absence of LPG indicated a weaker, or only occasionally modulated, connection to the other pyloric pacemaker neurons (PD and AB) (Figs. 7, 8). These data thus suggest that while IC activity strength is not affected by LPG photoinactivation, its ability to slow the pyloric rhythm requires the presence of LPG.

**Figure 7.**
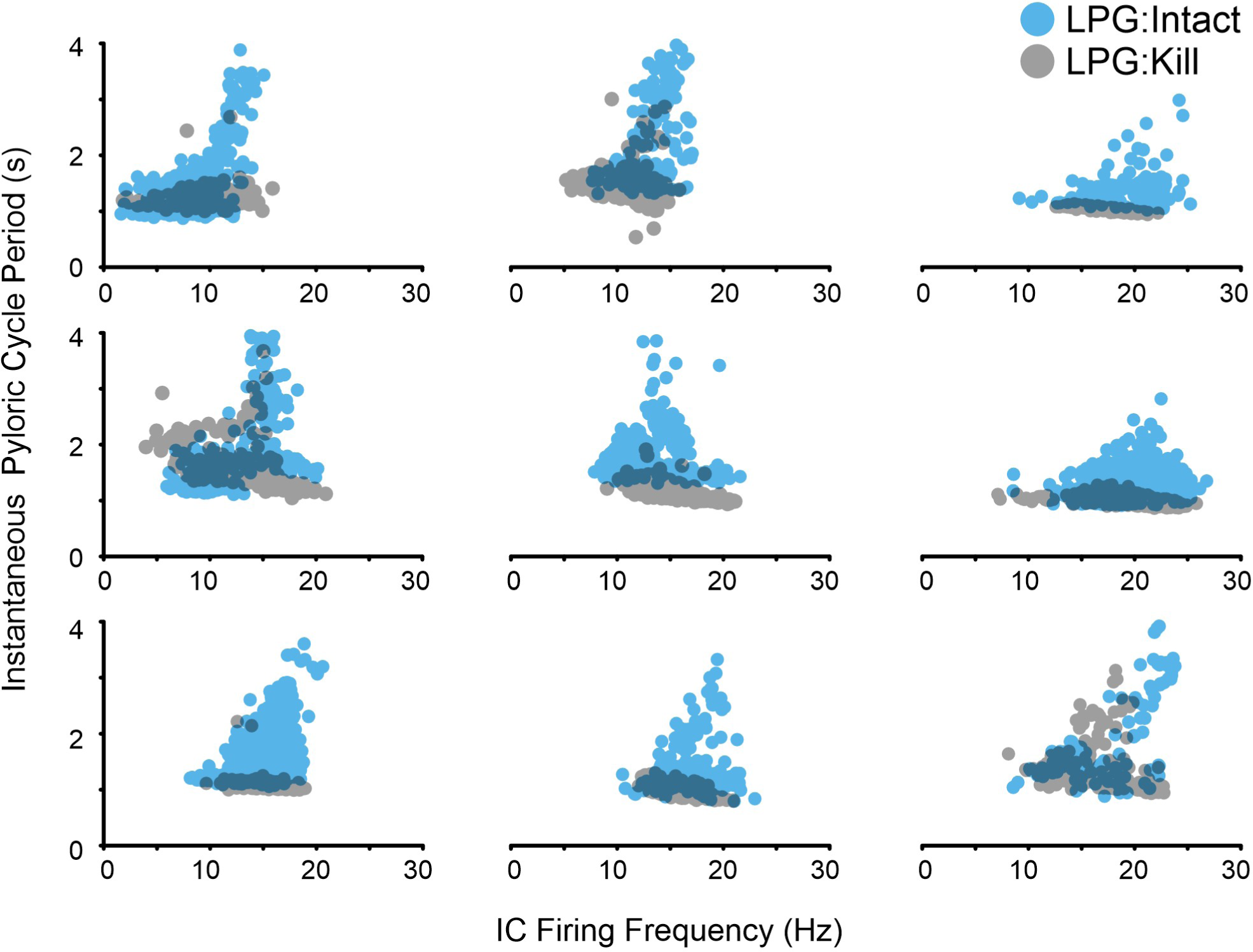
The same IC firing frequency did not alter pyloric cycle period in the absence of LPG neurons. Instantaneous pyloric cycle period is plotted against IC firing frequency in the LPG:Intact (blue dots) and LPG:Kill (grey dots) during 20 min windows. Each graph plots the data from a single experiment and each dot represents the instantaneous pyloric cycle period and the IC firing frequency within its pyloric-timed burst during the same pyloric cycle. Only cycle periods overlapping with IC gastric mill bursts (IC bursts > 0.45 s; see Fig. 6) are plotted.

**Figure 8.**
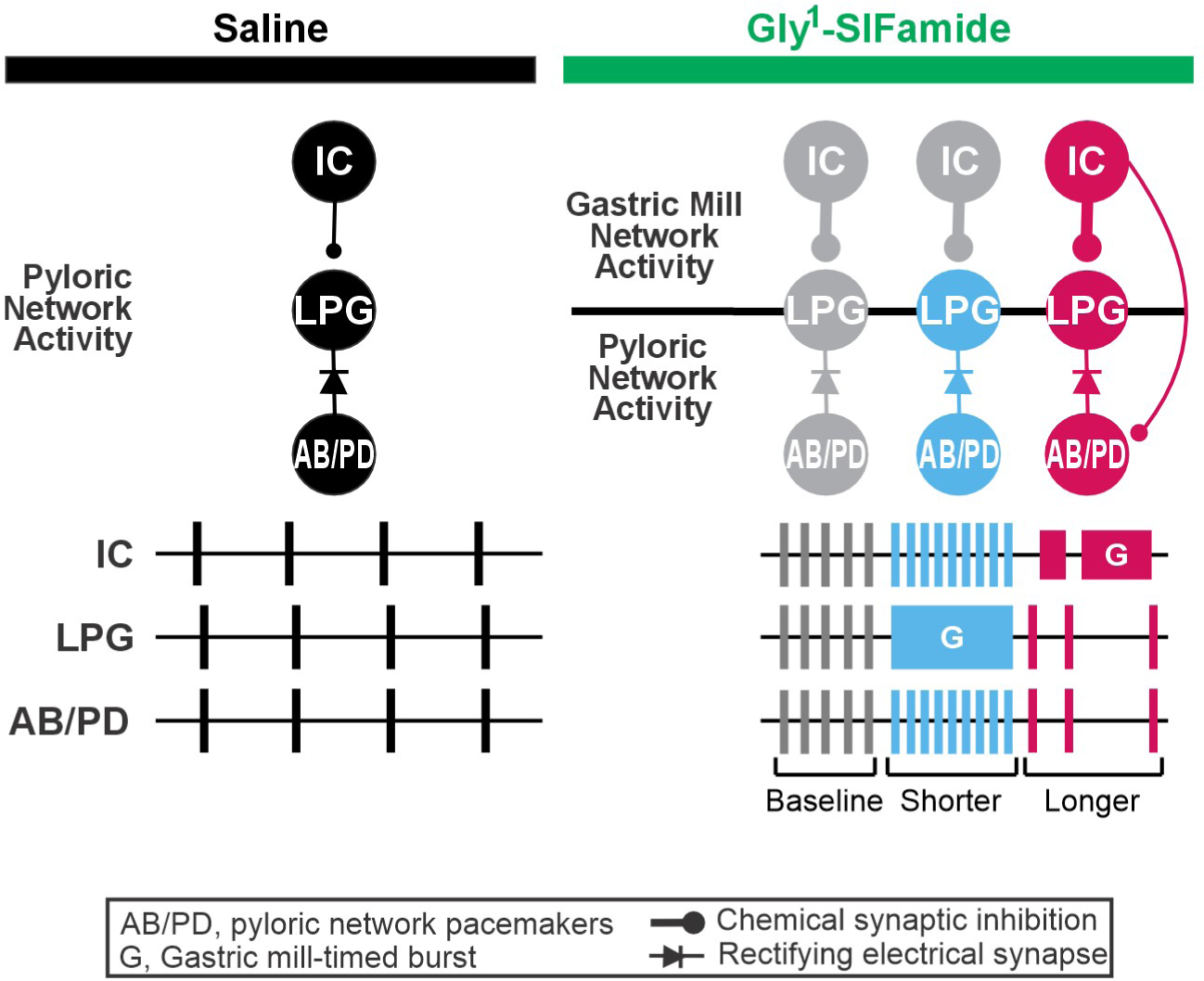
During the Gly^1^-SIFamide modulatory state, gastric mill bursting in LPG decreases pyloric cycle period, while gastric mill bursting in the IC neuron requires LPG to prolong the pyloric cycle period. In saline (*Left*, black traces and circuitry), LPG is active only in pyloric time, coincident with the AB/PD pyloric pacemaker neurons. Gly^1^-SIFamide (*Right*, green bar), elicits a unique gastric mill rhythm (Blitz et al., 2019; Fahoum and Blitz 2021, 2024), during which the pyloric cycle period varies between a baseline level due to Gly^1^-SIFamide modulation (grey), shorter periods which occur during LPG gastric mill-timed slow bursting (blue) and longer periods which occur during IC gastric mill bursting (pink). Gastric mill bursting in LPG (blue box, G: gastric mill burst) likely allows sufficient current though the rectifying electrical synapses (diode symbol) to the AB and PD neurons to decrease the cycle period of the pyloric rhythm (blue). IC gastric mill bursting (pink burst; G: gastric mill burst) is necessary for rhythmic increases in cycle period of the pyloric rhythm (pink). However, this inhibition appears to act primarily via IC chemical synaptic inhibition of LPG (thick pink stick/ball), with a possible weak contribution via chemical inhibition of the AB and/or PD neuron (thin pink line/ball). The IC to LPG inhibitory synapse is strengthened during Gly^1^-SIFamide modulation (compare thin line/ball in saline to thick line/ball in Gly^1^-SIFamide; Fahoum and Blitz, 2023).

## Discussion

In this study, we show that the cycle period of the faster pyloric rhythm varies between three levels in time with different phases of the related, slower gastric mill rhythm during the Gly^1^-SIFamide neuropeptide modulatory state. The LPG neuron switches from pyloric-only activity to dual pyloric and gastric mill activity during Gly^1^-SIFamide application (Fahoum and Blitz, 2021; Fahoum and Blitz, 2024; Snyder and Blitz, 2022). Surprisingly, we found that this dual-network LPG neuron is necessary for both phasic increases and phasic decreases in pyloric cycle period during this gastric mill rhythm version. Specifically, the pyloric cycle period decreases during the longer duration, intrinsically-generated gastric mill-timed bursts in LPG, and the pyloric cycle period is prolonged during IC neuron gastric mill bursts which coincide with a silent period in LPG. The complex biphasic regulation of a related network we identify here is distinct in both form and mechanism from previous examples of inter-network coordination in the STNS during other modulatory states, emphasizing the flexible nature of such coordination.

### Dual-network Neuron is Necessary for Biphasic Regulation

There are multiple ways in which coordination occurs between different behaviors and between their underlying CPG networks. Here we focused on a single modulatory state and how the activity of one network varies across a single cycle of a second network’s activity. Unlike other gastric mill rhythm versions, the Gly^1^-SIFamide gastric mill rhythm is triphasic (Blitz et al., 2019; Fahoum and Blitz, 2024). During this triphasic gastric mill rhythm, there is a “baseline” pyloric cycle period due to Gly^1^-SIFamide modulatory actions, and the pyloric rhythm is regulated away from this baseline in opposing directions during different phases of the slower gastric mill rhythm.

Photoinactivation to selectively eliminate the LPG neurons demonstrated that LPG is necessary for the phasic decreases in pyloric cycle period during the Gly^1^-SIFamide gastric mill rhythm. Decreased pyloric cycle periods occur during LPG gastric mill-timed bursts, which are intrinsically generated. In particular, Gly^1^-SIFamide enhances excitability and post-inhibitory rebound, and diminishes spike-frequency adaptation in LPG, enabling LPG to generate gastric mill-timed oscillations that do not require any synaptic input (Fahoum and Blitz, 2021; Snyder and Blitz, 2022). An LPG gastric mill-timed burst consists of a sustained depolarization during which LPG intrinsic currents apparently enable it to “escape” the influence of rhythmic hyperpolarizations occurring in its electrically coupled partners, the AB and PD neurons (Blitz et al., 2019; Fahoum and Blitz, 2021; Shruti et al., 2014; Snyder and Blitz, 2022). Electrical synapses are subject to modulation (Kothmann et al., 2009; Lane et al., 2018; O’Brien, 2014). Thus, it is possible that Gly^1^-SIFamide decreases the strength of electrical coupling between LPG and AB/PD to enable the periodic “escapes”. However, the electrical coupling apparently remains functional in Gly^1^-SIFamide, as AB/PD pyloric activity is necessary for LPG to generate pyloric bursting in this state (Fahoum and Blitz, 2021; Snyder and Blitz, 2022). Further, there is no apparent difference in coupling strength between saline and Gly^1^-SIFamide (Fahoum, Nadim, Blitz, unpublished). In addition to the continued presence of electrical coupling between LPG and AB/PD, we know that 1) decreases in pyloric cycle period require the presence of LPG (this study), 2) no other gastric mill neurons are active during LPG gastric mill bursts (Blitz et al., 2019; Fahoum and Blitz, 2024), and 3) LP and IC provide the only known intra-STG feedback to the pyloric pacemakers (Marder and Bucher, 2007; Nadim et al., 2011; Thirumalai et al., 2006; Zhao et al., 2011), but LP is photoinactivated or hyperpolarized and IC is silent or weakly active in pyloric time during this phase (Blitz et al., 2019; Fahoum and Blitz, 2024). Thus, although we are not able to selectively manipulate the electrical synapses between LPG and AB/PD to test our hypothesis, all available evidence indicates that the sustained depolarization during LPG gastric mill bursts has an excitatory effect on the AB/PD neurons to decrease the cycle period of the intrinsic pyloric bursting. This influence occurs despite rectification of the electrical coupling, which favors depolarizing current from AB/PD to LPG, but does allow for some current flow when LPG is depolarized relative to AB/PD (Shruti et al., 2014), which is apparently sufficient to decrease the pyloric cycle period.

Unexpectedly, photoinactivation of the LPG neurons also eliminated increases in pyloric cycle period which occur during IC or IC/LG bursting. With LPG intact, we found that LG promotes gastric mill-timed bursting in the IC neuron, but only IC, and not LG, is necessary for the extended pyloric cycle periods. The potential mechanisms for IC-elicited extension of the pyloric cycle period are that gastric mill-timed IC activity directly inhibits some or all of the pyloric pacemaker neurons, or IC gastric mill bursting excites another neuron that inhibits some or all of the pyloric pacemaker neurons. In the Gly^1^-SIFamide gastric mill rhythm, the only neuron besides LG that could be co-active with IC is the LP neuron, which could be “excited” via electrical coupling to IC and does provide inhibitory feedback to the pyloric pacemaker neurons (Nadim et al., 2011; Thirumalai et al., 2006; Weimann and Marder, 1994). However, in our experiments, to mimic the more physiological conditions of Gly^1^-SIFamide release from MCN5, with the coincident inhibition of LP via the MCN5 cotransmitter glutamate (Fahoum and Blitz, 2021), LP was always photoinactivated or hyperpolarized and therefore could not be inhibiting the pyloric pacemaker neurons. Thus, it is most likely that IC extends pyloric cycle period duration by inhibiting pyloric pacemaker neurons, which include 1 copy of AB, and 2 copies each of PD and LPG.

In many modulatory conditions, the AB neuron is a conditional oscillator and the PD and LPG neurons are active in pyloric time due to electrical coupling to AB (Ayali and Harris-Warrick, 1999; Marder and Bucher, 2007). In Gly^1^-SIFamide, LPG only generates pyloric-timed bursting if AB/PD do so, but AB/PD do not require LPG to generate pyloric-timed bursting (this study) (Fahoum and Blitz, 2024; Fahoum and Blitz, 2021; Snyder and Blitz, 2022). We found that selective photoinactivation of just the two LPG neurons almost entirely eliminated prolonged pyloric cycle periods. One possible explanation for this was that eliminating LPG impacted the activity of IC, however we found no change in IC gastric mill bursts. These results suggest that an IC to LPG synapse is primarily responsible for IC inhibition of the pyloric rhythm, effectively “funneling” IC chemical inhibition through LPG’s electrical coupling to the AB/PD neurons (Fig. 8). In fact, we recently determined that Gly^1^-SIFamide enhances a typically ineffective IC to LPG graded glutamatergic synapse, enabling gastric mill IC bursts to regulate LPG gastric mill-timed activity (Fahoum and Blitz, 2023). The current study indicates that the modulated IC to LPG synapse also indirectly regulates the pyloric cycle period via LPG’s electrical coupling to AB/PD. Some prolonged pyloric cycle periods in a subset of preparations when LPG was photoinactivated suggests an additional small contribution of an IC to AB and/or PD synapse, however it seems that the main IC regulation of pyloric cycle period occurs via inhibition of LPG (Fig. 8). The reason for IC chemical inhibition to act indirectly via the electrical synapse from LPG to AB/PD is not known. However, it may represent a conservation of function with the IC to LPG synapse playing roles in both gastric mill pattern generation as well as coordination between the pyloric and gastric mill networks.

Periodic long-duration IC bursts also inhibit the pyloric rhythm during activation of another modulatory input to the STG, the projection neuron MCN7, which releases the neuropeptide proctolin (Blitz et al., 1999). The specific neuron(s) targeted by IC to extend the pyloric cycle period in the MCN7 rhythm was not determined (Blitz et al., 1999). IC chemical synapses within the STG, including the IC to LPG synapse, are fast inhibitory glutamatergic connections that are blocked by the chloride channel blocker picrotoxin (Fahoum and Blitz, 2024; Marder and Bucher, 2007). Inhibitory neuron connections between networks are implicated in other inter-network interactions, although further exploration is still needed for a mechanistic understanding at the cellular level (Huff et al., 2022).

### Diverse Mechanisms of Inter-network Coordination

Distinct mechanisms have been identified for inter-network coordination in the STNS and other systems. In the STNS, other gastric mill rhythm versions are biphasic, and when pyloric rhythm variability has been quantified, the pyloric period during one phase of the gastric mill rhythm is at a “baseline” period due to modulatory input, and regulated away from this baseline during the other phase of the gastric mill rhythm (Bartos and Nusbaum, 1997). For instance, during a gastric mill rhythm elicited by the modulatory neuron MCN1, the LG neuron presynaptically inhibits MCN1 at its entrance into the STG, rhythmically decreasing modulatory excitation of the pyloric network during the LG phase (Bartos and Nusbaum, 1997; Coleman et al., 1995; Coleman and Nusbaum, 1994; Nusbaum et al., 1992). Thus, there is indirect communication between the two networks via a local feedback synapse. Indirect interactions between networks can also occur via long-distance feedback from network neurons to modulatory inputs or via motor efference copy to related networks (Blitz and Nusbaum, 2008; Lambert et al., 2023; Wood et al., 2004). Although there are both local and long-distance feedback projections to the source of Gly^1^-SIFamide (MCN5) (Blitz et al., 2019; Norris et al., 1996), they do not contribute to the inter-network regulation described here, as the neuropeptide was bath-applied. Instead, we found that synapses between network neurons underlie biphasic regulation of the pyloric rhythm.

In addition to phasic coordination, there may be correlated changes in the activity levels of related networks such as increased respiratory activity to match more intense forms of locomotion, or increased swallowing occurrences to match increased chewing rates (Hao et al., 2021; Hérent et al., 2023). When the pyloric and gastric mill networks are both active, their activity level is coordinated such that faster pyloric rhythms occur with faster gastric mill rhythms (Bartos et al., 1999; Powell et al., 2021; Saideman et al., 2007). Correlated changes in timing and/or strength can reflect common inputs. In the crab STNS, the modulatory neuron MCN1 acts on both networks, and increasing MCN1 firing frequency speeds both rhythms (Bartos et al., 1999; Bartos and Nusbaum, 1997). Similarly, in lamprey, rodents, and cats, the mesencephalic locomotor region provides parallel input to respiratory and locomotor CPGs (Hérent et al., 2023; Juvin et al., 2022; Ryczko and Dubuc, 2013).

Some related behaviors display coupling of their activity, with a discrete number of cycles of one pattern occurring relative to a full cycle of the other pattern. Coupling occurs between locomotion and chewing, and locomotion and respiration in humans, other mammals and birds; between whisking and respiration in rodents; and between pyloric and gastric mill rhythms in crustaceans (Bartos et al., 1999; Clemens et al., 1998a; Juvin et al., 2022; Maezawa et al., 2020; Moore et al., 2013; Powell et al., 2021). Coupling of rhythmic behaviors may arise from mechanical forces. For example, flight muscle attachment to structures essential for respiration in birds, and the movement of internal organs in galloping horses may contribute to coupling between locomotion and respiration (Juvin et al., 2022). Similarly, accessory teeth movements are mechanically coupled to the cardiopyloric valve in the stomatogastric system (McGaw and Curtis, 2013). However, direct synaptic connections between CPGs can also contribute to this type of inter-network coordination (Juvin et al., 2022). In rodents, proper timing of swallowing within the respiratory cycle, and 1:1 coupling of whisking and fast sniffing, appear to be due to synapses between these related CPGs (Huff et al., 2022; Moore et al., 2013). These coupling ratios between networks are not fixed and can vary during different forms of locomotion, or different breathing behaviors such as fast sniffing and slow breathing (Juvin et al., 2022; Moore et al., 2013; Saunders et al., 2004). In the crab STNS, a direct inhibitory synapse from the pyloric pacemaker neuron AB onto the gastric mill CPG neuron Int1 is essential for coupling between the pyloric and gastric mill rhythms driven by the modulatory neuron MCN1. This inter-network synapse allows the pyloric rhythm to control the speed of the gastric mill rhythm (Bartos et al., 1999; Nadim et al., 1998). However, during the Gly^1^-SIFamide rhythm, the speed of the pyloric rhythm does not regulate LPG gastric mill bursting, reinforcing the flexible nature of inter-network coordination (Fahoum and Blitz, 2024).

Our results here suggest that the same rectifying electrical synapse alternately reinforces and diminishes modulatory neuropeptide effects on network output. These actions occur due to intrinsic currents enabling reinforcement, and chemical synaptic currents leading to diminished peptide actions. LPG thus serves as a point of convergence for coordination, despite the two network interactions acting in opposite directions. Due to the fluid nature of network composition, dual-network neurons such as LPG provide flexibility by adding and subtracting their activity to different behavioral outputs (Ainsworth et al., 2011; Clapp et al., 2011; Dickinson et al., 1990; Hooper and Moulins, 1989; Roopun et al., 2008; Weimann et al., 1991). Additionally, dual-network neurons can provide flexibility in coordinating network activity by serving as a variable link between networks. This ability however depends on whether they are recruited into new networks as passive followers, or as active components with sufficient synaptic access to a new network to mediate coordination (Fahoum and Blitz, 2024; Hooper and Moulins, 1990).

Dysfunctional inter-network coordination can impact health and well-being, such as an insufficient oxygen supply relative to metabolic demand if respiration is not coordinated with other motor activity, decreased food intake or choking due to improper coordination of breathing and swallowing, or disrupted communication abilities arising from problems coordinating vocalization and respiration (Barlow, 2009; Juvin et al., 2022; Yagi et al., 2017). Thus, it is important to continue adding to our understanding of inter-network coordination among the many behaviors that require coordination, including understanding how coordination varies across behavioral and environmental conditions (Clemens et al., 1998b; Juvin et al., 2022; Stein and Harzsch, 2021; Stickford and Stickford, 2014).

## Contributions

S-RHF and DMB designed research, S-RHF and BG performed research, BG analyzed data, DMB and BG wrote the paper, DMB, S-RHF, and BG edited the paper

## Acknowledgements

We thank Michael Hughes and Anh Vo (Statistical Consulting Center, Miami University) for assistance with statistical analysis.

## Conflict of Interest

The authors report no conflict of interest.

## Funding sources

National Science Foundation IOS:1755283 (DMB), Biology Department, Miami University

## Notes

### Competing Interest Statement

The authors have declared no competing interest.

### Summary of Updates

Title, Discussion section, and Figure 8 revised.

